# Nonlinear Parameter and State Estimation Approach for Intradialytic Measurement of Absolute Blood Volume

**DOI:** 10.1101/2023.08.12.553092

**Authors:** Rammah Abohtyra, Tyrone Vincent, Daniel Schneditz

**Author notes:** Corresponding author: Rammah Abohtyra,.

## Abstract

**Background:** Managing blood and fluid volumes in chronic kidney disease (CKD) patients plays an essential role in dialysis therapy to replace kidney function.

**Objective:** This study aims to develop an estimation approach to provide predictable information on blood and fluid volumes during a regular dialysis routine.

**Methods:** The method utilizes a non-linear fluid volume model, an optimization technique, and the Unscented Kalman Filter (UKF). This method does not rely on specific ultrafiltration and dilution protocols and uses the Fisher information matrix to quantify the estimation error.

**Results:** The method was applied to 21 data sets of ten patients. A significant moderate correlation was obtained when estimated blood volumes were compared to a different method applied to the same data set. Average specific blood volumes were plausible and in the range of 78.7 and 75.9 mL/kg at the end of the high ultrafiltration rate pulse and above the critical level of 65 mL/kg. Critical blood volumes were only observed in four studies done on three patients.

**Conclusion:** The absolute blood volume estimated at the beginning and during every dialysis session offers the opportunity to detect critical blood volumes and to improve fluid management in CKD patients significantly.

## 1 Introduction

Chronic kidney disease (CKD) is a major health concern, significantly decreasing the patient’s quality of life, causing a high mortality, and a substantial economic impact [1]. Most CKD patients receive an intermittent therapy of 3-4 hemodialysis sessions (each session is 2-5 h long) per week to support or to completely replace their lost kidney function. Apart from water soluble metabolites removed by diffusion and adjustment of acid-base and electrolyte balance when blood is exposed to dialysate in the extracorporeal circulation, excess fluid accumulating in the typical 2-day interval throughout the body and leading to an intradialytic weight gain in the range of 2-4 kg has to be removed by ultrafiltration (UF) of blood to re-establish so-called dry weight at the end of each single treatment session [2].

Intradialytic hypotension (IDH) is a common complication of hemodialysis and is associated with a higher risk of cardiovascular morbidity and mortality [3–6]. The single most important factor for IDH is acute hypovolemia, either because of errors in prescribing the proper ultrafiltration volume as the true dry weight is essentially unknown, or because of prescribing ultrafiltration rates largely exceeding the rates at which extravascular fluid is refilled into the vascular compartment. Depending on the patient’s hemostatic defense mechanisms to compensate for acute hypovolemia, IDH may then develop with all unwanted side effects. Therefore, knowing a patient’s blood and fluid volumes at the start of dialysis and during ultrafiltration could allow clinicians to avoid critical degrees of hypovolemia and to return CKD patients to their dry weight safely [7].

Dry weight, adequate ultrafiltration, and absolute blood volume have been of concern since the early days of dialysis. However, most standard measuring methods are not feasible in everyday clinical practice so that ultrafiltration of excess volume remains largely prescribed by clinical judgment only.

All methods to measure blood volume are based on some form of indicator dilution and require the precise administration and measurement of that indicator. The removal of excess fluid by ultrafiltration itself can also be seen as a form or indicator dilution, where fluid volume is removed (rather than added) and the indicator, in this case red blood cells is therefore concentrated (rather than diluted). Several techniques for on-line measurement of blood components such as red blood cells are now available in clinical practice, and the question arises whether they can be used for absolute volume assessment in clinical routine.

This work therefore aims to estimate blood volume during dialysis, to predict the change in blood volume during UF, and to detect a low absolute blood volume by monitoring the concentration of red blood cells recorded during routine treatments in response to ultrafiltration without relying on a specific ultrafiltration or volume infusion protocol. Instead, the method utilizes dialysis data generated in a regular dialysis routine using standard UF profiles.

## 2 Materials and Methods

### 2.1 Mathematical Model

The volume kinetics during ultrafiltration is described by a previously published two-compartment extracellular fluid volume model [8, 9]. Some modifications such as a constant protein mass in each compartment and a lymphatic flow component were introduced to the model [8, 10–12] (Fig. 1). The states of the model are the plasma volume, *V*_*pl*_ (L), and the interstitial fluid volume, *V*_*int*_ (L); while the input, *Q*_*u*_, is the ultrafiltration rate (UFR) (L/min). The output is the whole body hematocrit (*H*), which is the volume fraction of all red blood cells (*V*_*rbc*_, L) in total blood volume (*V*_*b*_ = *V*_*rbc*_ + *V*_*pl*_), i.e., *H* = *V*_*rbc*_*/V*_*b*_.

**Fig. 1.**
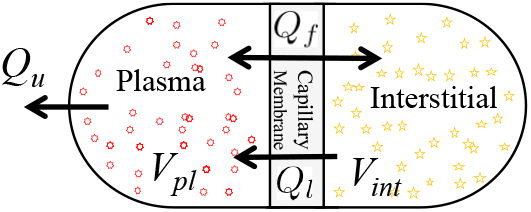
The two-compartment model consists of interstitial fluid volume (*V*_*int*_) and intravascular (plasma) fluid volume (*V*_*pl*_) separated by a microvascular membrane; arrows indicate microvascular (*Q*_*f*_), lymph (*Q*_*l*_), and ultrafiltration flows (*Q*_*u*_).

The mathematical description of this model during dialysis is given by the nonlinear differential system:

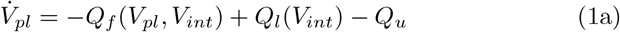

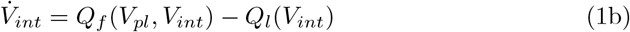

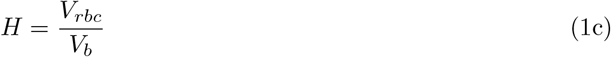

where *Q*_*f*_ (L/min) represents the microvascular (shift/refill) filtration flow, in which, *K*_*f*_ (L/min.mmHg) is the transcapillary permeability coefficient; and *Q*_*ly*_ (L/min) represents the lymphatic flow. All equations are provided in the supplementary file.

Additionally, the absolute blood volume *V*_*b*_(*t*) is given as the sum of variable plasma volume *V*_*pl*_(*t*) and constant red blood cell volume *V*_*rbc*_. Six parameters including initial conditions for plasma (*V*_*pl*_(0)) and interstitial volumes (*V*_*int*_(0)) are considered patient specific *θ* = [*V*_*pl*_(0), *V*_*int*_(0), *V*_*rbc*_, *K*_*f*_, *d*_1_, *d*_2_]^*T*^, whereas the other parameters in Table 1 (supplementary file) are assumed nominal for all patients.

**Table 1.** Glossary of terms.

| Symbol | Generic term |
| --- | --- |
| $V$ | Volume (L) |
| $v$ | Specific blood volume (mL/kg) |
| $M$ | Mass (kg) |
| $C$ | Concentration (mg/L) |
| $Q$ | Fluid flow rates (volume/time) |
| UFR, ($Q_u$) | Ultrafiltration rate (volume/time) |
| $V_u$ | Ultrafiltration volume (L) |
| $t$ | time (min) |
| $T_s$ | Sampling time (min) |
| $E$ | Expected value |
| $H$ | Hematocrit (%) |
| $pl, int, e$ | subscript, referring to plasma, interstitial, extracellular |
| $b, rbc, f$ | subscript, referring to blood, red blood cell, filtration |
| $l, c, b$ | subscript, referring to lymph, capillary, blood |
| $\theta, \hat{\theta}$ | referring to parameters, estimated parameters |
| $g, h, \beta$ | constants |
| $p$ | Hydrostatic pressure (mmHg) |
| $\pi$ | Colloid-osmotic pressure (mmHg) |

### 2.2 Parameter Estimation

The estimation approach utilizes a general nonlinear model of a continuous-time differential equation described by a state transition (*f*_*s*_) and observation function (*h*_*o*_):

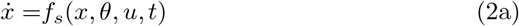

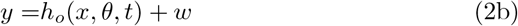

where *x* ∈ ℝ^*n*^ denotes the state of the system; *u* denotes the input to the system; and *θ* ∈ ℝ^*m*^ contains the model parameters whose values (or some of them) are estimated from data; *y* is the model output; *t* is the time *t*_0_ ≤ *t* ≤ *t*_*final*_ between initial, *t*_0_, and final time, *t*_*final*_; and *w* represents measurement noise.

The algorithm optimizes the parameters *θ* by minimizing the sum of the square errors between the measured (*H*_*i*_) and the model’s (*y* (*θ, t*_*i*_)) (1) in response to the ultrafiltration (*u*) as an input to the model, where *t*_*i*_ indicates the time of *H* measurements (*i* = 1, …, *N*). The objective function is given by

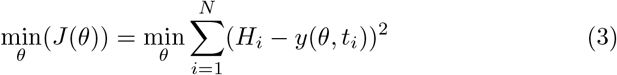

where *H*_*i*_ and *y* (*θ, t*_*i*_) refer to observed measurements and output trajectory of the nonlinear model, respectively. The optimal solution of (3) is denoted by 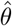. The estimation algorithm consists of two nested loops (Fig. 2): the outer one loops over a random set of initial conditions *θ*_*n*,0_, and the inner loop is based on the Levenberg-Marquardt method where the value *θ*_*n*_ is updated on the *i*th cycle by *θ*_*n,i*+1_ = *θ*_*n,i*_ − ∇_*n,i*_ where ∇_*n,i*_ uses the steepest descent method.

**Fig. 2.**
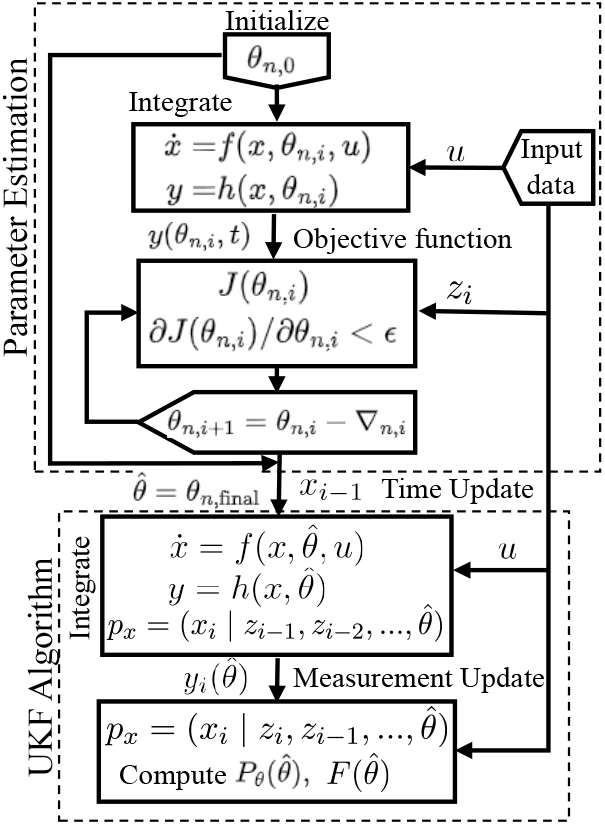
Schematic representation for algorithm 1. The parameter estimation and Unscented Kalman Filter (UKF) prediction algorithms are independently performed. The estimated parameter 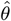 is an input to the UKF algorithm.

Note that the estimated parameter 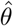 is an input to the Unscented Kalman Filter (UKF) prediction algorithm.

### 2.3 Prediction

The UKF [13, 14] is integrated with the nonlinear model and data to compute the estimation error using the Fisher information matrix, to update states (blood and fluid volumes), and to predict the measured hematocrit.

To estimate the parameter uncertainty and make predictions of state trajectories, using UKF, we introduce the model uncertainty (*n*) into (2a)

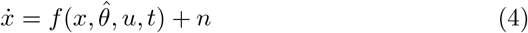

and assume that the distribution of model uncertainty (*n*) and measurement noise (*w*) in (2b) is piece-wise constant over intervals of *T*_*s*_ (*T*_*s*_ is a sampling time), the magnitude for the interval *k* (*k* is an index) is a Gaussian random vector with zero mean given by

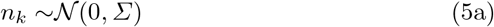

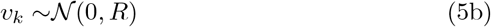

and where 𝒩 represents the notation for normal Gaussian distribution; *Σ* and *R* are the covariance of model uncertainty and measurement noise, respectively. Consequently, we assume *x* is Gaussian. Since *y* is conditionally affected by the state *x*, we also assume that *y* is Gaussian. Thus, the conditional state and output density functions are Gaussian:

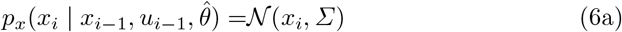

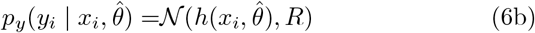

where *x*_*i*_ is the solution to (2) with *n* = 0, initial condition *x*_*i* −1_ and input *u*_*i*−1_.

As illustrated in Fig. 2, the prediction algorithm uses the UKF with the estimated parameter 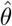 and the nonlinear model (2) to compute the Fisher information matrix 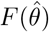, update states, and predict the output of this model. Here, we provide iterative steps for updating the UKF, utilizing observations *z*_0_, *z*_1_, …, *z*_*N*_ at a sampling time *T*_*s*_, and an input *u*, where *u* is held as a piecewise constant during the interval (*i* − 1)*T*_*s*_ ≤ *t* ≤ *iT*_*s*_:

- 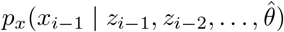 represents the previous estimated state.
- 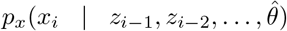 represents the prediction update step (time update), obtained by integrating model (2) between time *t* = (*i* − 1)*T*_*s*_ to *t* = *iT*_*s*_, with initial conditions 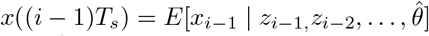.
- 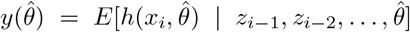 represents the prediction step of a measurement *z*_*i*_.
- 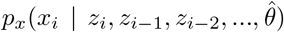 represents the measurement update step of the UKF.

Algorithm 1 below presents the parameter estimation (A) and prediction (UKF) algorithm (B).

### 2.4 Parameter Estimation Sensitivity Analysis

We use the Fisher information matrix [15] to quantify the quality of the estimation based on model uncertainty and measurement noise. This analysis uses the following assumptions.

- The distribution of *y* can be approximated by a Gaussian distribution.
- The experimental uncertainty is captured by model (2)
- The covariance of the prediction error is independent of *θ*.
- The UKF is approximately optimal, so that the prediction errors are uncorrelated.
- The parameter estimates are unbiased

#### Theorem 1

Under the assumptions listed above, the covariance of the parameter estimates (P_θ_) is bounded below by

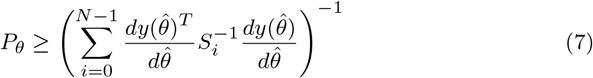

*where S*_*i*_ *is the covariance of the prediction error calculated using the UKF*.

#### Algorithm 1

The Parameter and Prediction Algorithms

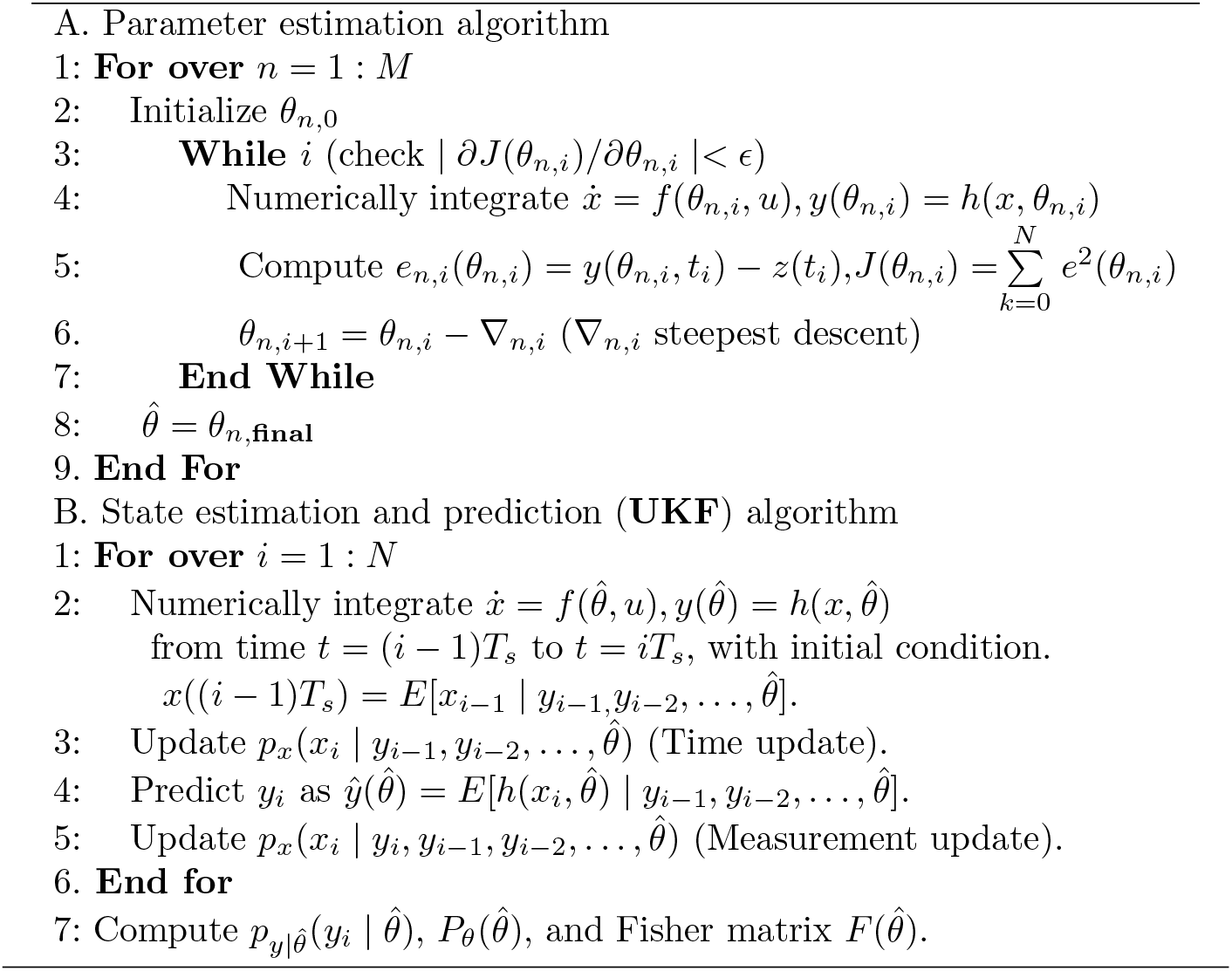

**Proof:** For unbiased parameter estimates, the parameter estimate covariance is bounded below by the inverse of the Fisher information matrix (see e.g. [15]), which is calculated from the likelihood function. The likelihood function for the measurements *Z* where *Z* = [*z*_0_ · · · *z*_*N*_]^*T*^ is given by

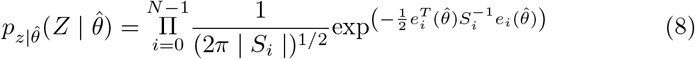

where 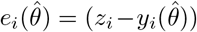, in which 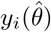 is the UKF one step ahead prediction; and *S*_*i*_ = *GP*_*i*_*G*^*T*^ + *R* is the covariance of the error 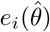, where *P*_*i*_ is the estimated state error covariance at time *i*, and *H* is the mapping from state to output (i.e., *G* = [ 1 0 ] for the model (1)). Then the log-likelihood function is

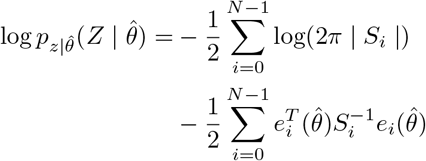

Then

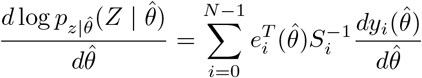

The Fisher information matrix is given by the covariance of this expression, so we find

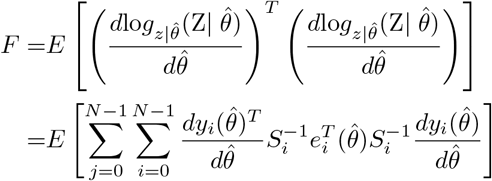

Under the assumption that the prediction error 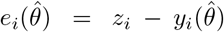 is uncorrelated,

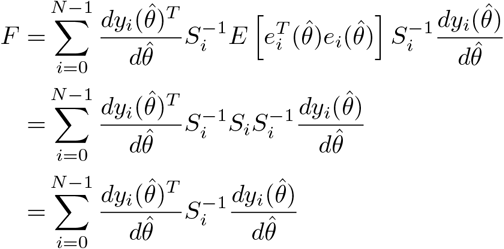

The result follows by inverting the Fisher information matrix. □

### 2.5 Initial Conditions

Initial conditions were based on physiologic considerations. Normal blood is about 70 mL/kg and scales with body mass [16]. The ratio of blood to extracellular volume is about 1/3 [17]. This ratio is maintained with moderate volume expansion [18]. Volume expansion was determined from the difference of pre-to post-treatment body mass. Interstitial protein concentration is about 1/3 of plasma protein concentration. The protein mass (*M* = *C* ∗ *V*) in each of the compartments was assumed as constant.

- Initial blood volume *V*_*b*_(0) = 70 mL/kg * weight post + (weight pre -weight post)/3.
- Initial extracellular volume *V*_*e*_(0) = (3*70 mL/kg)* weight post+(weight pre-weight post).
- Initial plasma volume *V*_*pl*_(0) = *V*_*b*_(0) ∗ (1− *H*(0)).
- Initial interstitial volume *V*_*int*_(0) = *V*_*e*_(0) − *V*_*pl*_(0).
- Initial microvascular coefficient *K*_*f*_ = 8.1e-5 (L/(min.mmHg.kg))*weight post [8].
- Initial plasma protein mass *M*_*pl*_(0) = *C*_*pl*_(0) ∗ *V*_*pl*_(0).
- Initial interstitial protein mass *M*_*int*_(0) = (*C*_*pl*_(0)*/*3) ∗ *V*_*int*_(0).
- Red blood cell volume *V*_*rbc*_ = *V*_*b*_(0) ∗ *H*(0).

These relationships create a patient specific initial condition *θ*_0_ = [*V*_*pl*_(0), *V*_*int*_(0), *V*_*rbc*_, *K*_*f*_, *d*_1_, *d*_2_]. Upper and lower boundaries of *θ*_0_ within plausible physiological ranges (±50% of *θ*_0_) centered at *θ*_0_ create randomly initial conditions Θ_0_ for each patient. To address the sensitivity to initial conditions (initial guess for *θ*), the uncertainty and uniqueness of the optimal solution was examined using randomly selected sets of initial parameters Θ_0_ (positive and uniform distribution) within bounded intervals. The output provides a distribution of optimal solutions represented by Kernel Probability Density (KPD) functions estimated over the random initial values Θ_0_ and describes the quality of a given estimation.

### 2.6 Clinical Data

The method was tested with hematocrit and ultrafiltration data collected in a previous study approved by the local internal review board. In this previous study a different method to estimate absolute blood volume in hemodialysis patients using ultrafiltration pulses at two time points (*v*_1_, *v*_2_) was examined [19]. Patient information such as sex, body mass at treatment end, ultrafiltration volume, and initial plasma protein concentration were also available and used for parameter estimation (Table 2, supplementary file). Treatments are identified by a treatment ID where the leading digits refer to the patient, and where the last digit refers to the measurement done on different treatment days.

**Table 2.** Estimation results (mean*±*SD) of blood and fluid volumes for 10 patients (P1 to P10) with 21 treatments, initial blood volume *V*_*b*_(0), the minimum specific blood volume *v*_1_ and *v*_2_ at UF steps, and initial plasma and interstitial volumes *V*_*pl*_(0), *V*_*int*_(0). The last digit of the treatment ID refers to the measurement in the same patient done on different treatment days, * specific blood volume ≤ 65 mL/kg

| Patient ID | $V_b(0)$<br>L | $v_1$<br>mL/kg | $v_2$<br>mL/kg | $V_{pl}$<br>(L) | $V_{int}$<br>(L) | MSE<br>(-) |
| --- | --- | --- | --- | --- | --- | --- |
| P11* | 6±0.25 | 69.4 | 62.6 | 4.3±0.18 | 15±1.18 | 3.1e-4 |
| P12* | 5.4±0.07 | 63.9 | 60.0 | 3.9±0.05 | 14.7±1.2 | 2.6e-4 |
| P21 | 5.1±0.50 | 75.1 | 73.1 | 4.1±0.4 | 17.1±3.13 | 3.8e-6 |
| P22 | 5.4±0.52 | 81.3 | 81.0 | 4.4±0.42 | 20.5±3.84 | 6.9e-5 |
| P23 | 5.5±0.28 | 82.6 | 82.2 | 4.5±0.23 | 20.7±2.02 | 6.9e-5 |
| P31 | 5.1±0.35 | 85.5 | 85.1 | 3.8±0.26 | 14.5±2.8 | 6.3e-5 |
| P32 | 5.2±0.85 | 88.1 | 81.3 | 3.9±0.43 | 16.2±2.65 | 2.6e-4 |
| P33 | 5.7±0.85 | 98.3 | 92.5 | 4.3±0.64 | 16.5±4.24 | 5.7e-5 |
| P41 | 5.2±0.63 | 73.7 | 72.9 | 3.9±0.47 | 16.7±4.25 | 2e-4 |
| P42 | 6.5±0.40 | 95.1 | 97.4 | 4.8±0.30 | 27.2±2.99 | 6.3e-5 |
| P43 | 5.9±0.21 | 85.3 | 85.5 | 4.4±0.15 | 24.6±1.66 | 4.8e-5 |
| P51 | 5.8±0.16 | 95.4 | 94.3 | 4.6±0.13 | 27.9±1.7 | 1.3e-5 |
| P52 | 5.7±0.28 | 89.2 | 90.9 | 4.6±0.23 | 22.2±2.11 | 3.2e-4 |
| P61 | 5.5±0.84 | 88.5 | 79.1 | 4.1±0.64 | 16.4±4.12 | 5e-5 |
| P62 | 5.1±0.78 | 76.1 | 67.5 | 3.6±0.58 | 13.9±3.76 | 1e-4 |
| P71* | 5.1±0.64 | 42.1 | 41.6 | 4.0±0.49 | 17.5±4.27 | 8.7e-5 |
| P81 | 5.3±0.73 | 72.8 | 69.7 | 4.0±0.56 | 15.7±3.46 | 4.9e-5 |
| P82 | 5.6±0.91 | 77.3 | 75.7 | 4.3±0.69 | 18.3±4.97 | 6.6e-5 |
| P83 | 5.2±1.14 | 72.4 | 70.1 | 3.9±0.87 | 15.8±6.19 | 1.3e-4 |
| P91 | 4.7±0.35 | 80.4 | 74.9 | 3.7±0.28 | 14.8 ±2.26 | 8.1e-5 |
| P101* | 4.3±0.58 | 60.9 | 56.7 | 3.4±0.45 | 13.3 ±2.46 | 2e-4 |
| Average | 5.4±53 | 78.7 | 75.9 | 4.12±0.4 | 18.1 ±3.1 | 1.2e-4 |

We used a 60 min training phase of combined ultrafiltration rate input and hematocrit data to train the model and to estimate the parameter *θ*, and then the rest of the data was used to validate the results and test the quality of the model.

### 2.7 Statistics

Data are presented as average values ± standard deviation (SD) or coefficients of variation (CV) unless otherwise indicated. Variables obtained by different methods were compared by linear regression and Bland-Altman analysis.

The quality of the output was quantified by a normalized mean square error (MSE) between modeled and measured variables:

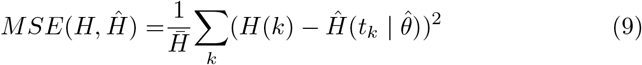

where 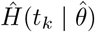 is the modeled hematocrit at time interval *k* of the measured hematocrit taken from the model’s output trajectory generated by integrating the model forward with the estimated parameters, and 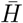 is an average value of the measured hematocrit.

## 3 Results

The time course of hematocrit measured in 21 hemodialysis treatments was successfully modeled by the new algorithm and provided plausible model parameters (Table 2 and Table 3 supplementary file). A representative example of the typical time course of measured, modeled, and predicted hematocrit is shown in Fig. 3. As expected, hematocrit increased when fluid volume was removed from plasma volume by ultrafiltration, and dropped when ultrafiltration was stopped and fluid was refilled from the expanded interstitial space. Overall, there was an increase in hematocrit with ongoing ultrafiltration, and the increase was smooth with constant ultrafiltration rate.

**Fig. 3.**
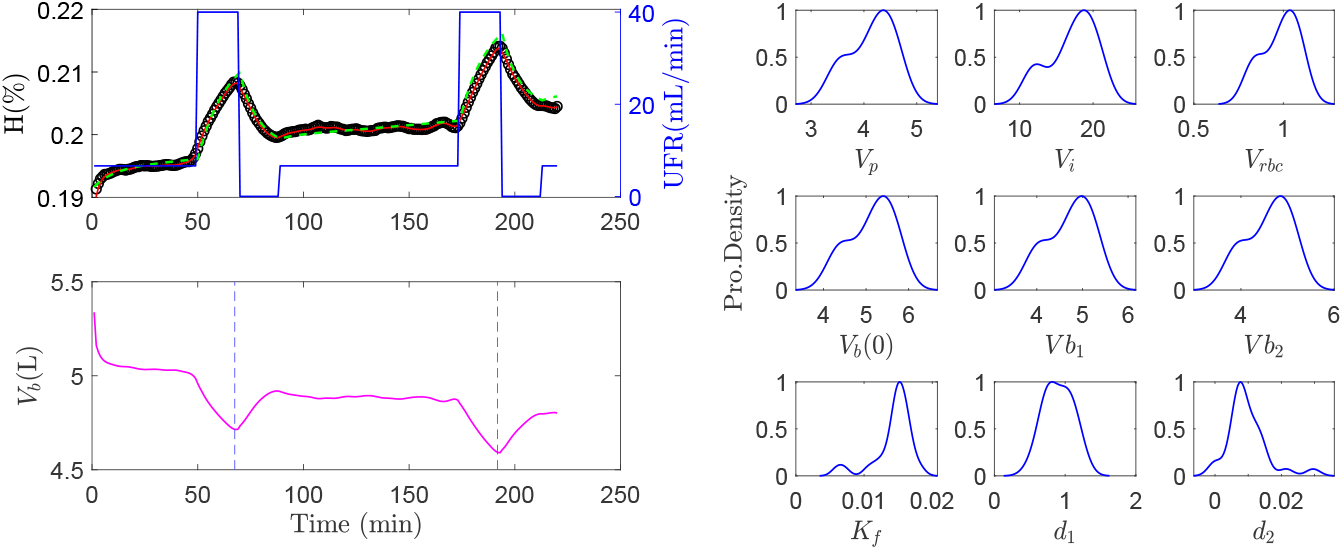
Modeling and prediction. Left, top panel: Modeled hematocrit (green dashed trajectory) compared to predicted hematocrit (red trajectory) using UKF versus actual hematocrit (black circles) (data from treatment P21) in response to variable ultrafiltration rate (UFR, blue line). Left, bottom panel: Predicted blood volume (magenta) and time points (dashed blue lines) of blood volumes estimated at the end of ultrafiltration steps (*v*_1_, *v*_2_). Right panels: KPD functions for various parameters over the random initial solution Θ_0_ estimated for treatment P21.

Figure 3 shows an example of a well-modeled hematocrit. The hematocrit prediction using the UKF approach is only unremarkably better than the modeled hematocrit. The KPD functions show unimodal and close to normal distributions around the estimated maxima.

A summary plot of 20 individual measurements is shown in Fig. 4. The corresponding model parameters of all 21 studies are listed in Tables 2 and Table 3 (supplementary file).

**Fig. 4.**
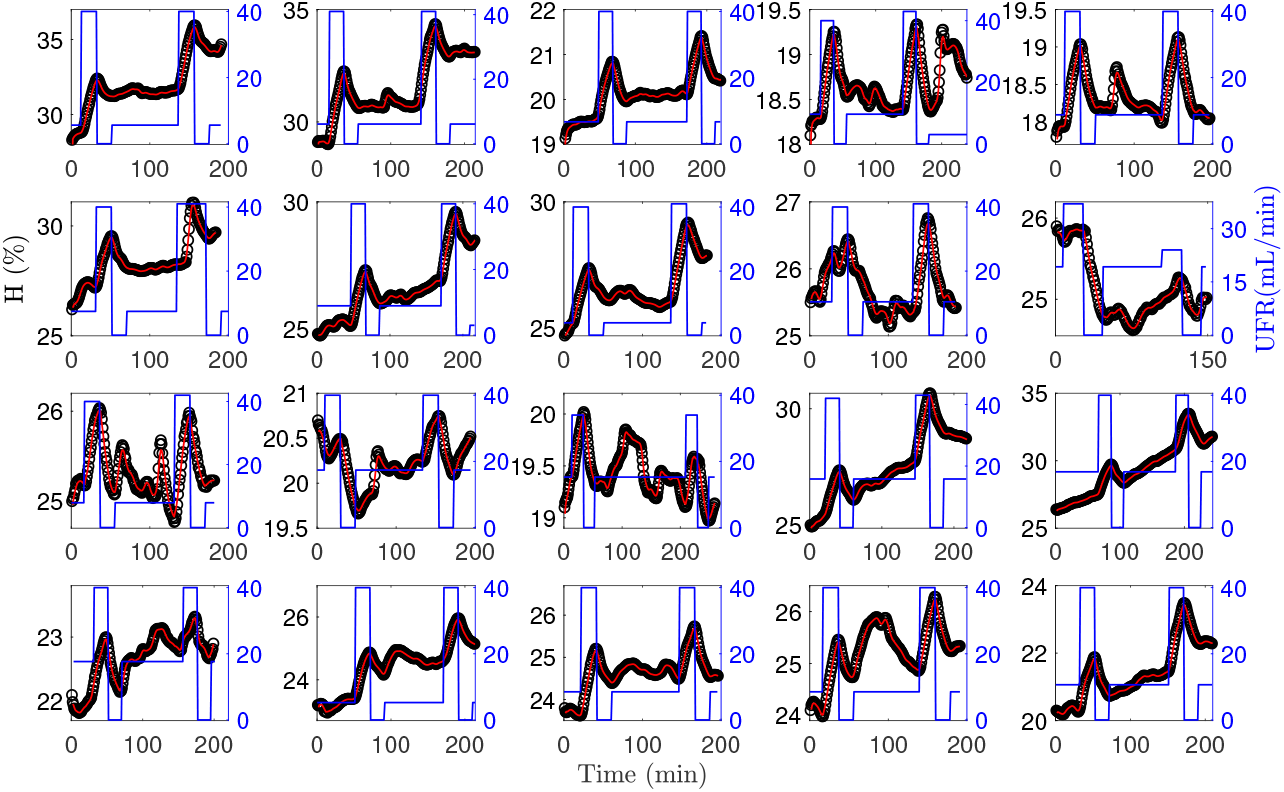
Prediction and measurement. Comparison of predicted (red lines) and measured (black circles) hematocrit in 20 treatments done in 10 patients using ultrafiltration rate profiles (blue lines) (one measurement is omitted because of limited space)

Notice that all parameters of interest are obtained with distinct standard deviations (Table 2). The average CV for estimated blood and plasma volumes was in the range of 10%. Specific blood volumes were plausible and in the range of 78.7 and 75.9 mL/kg at the end of the high ultrafiltration rate pulse (Table 2). Specific volumes below the critical level of 65 mL/kg were observed in four studies done in three patients, in one of those (P71), absolute blood volume was already below the critical at treatment start.

A comparison of volumes obtained by the new approach with results from the previous analysis is provided in Fig. 5. The correlation (*r*) is significant, albeit moderate (0.5 ≤ *r* ≤ 0.85). The old method overestimates blood volumes in the high-volume range when compared to the new method.

**Fig. 5.**
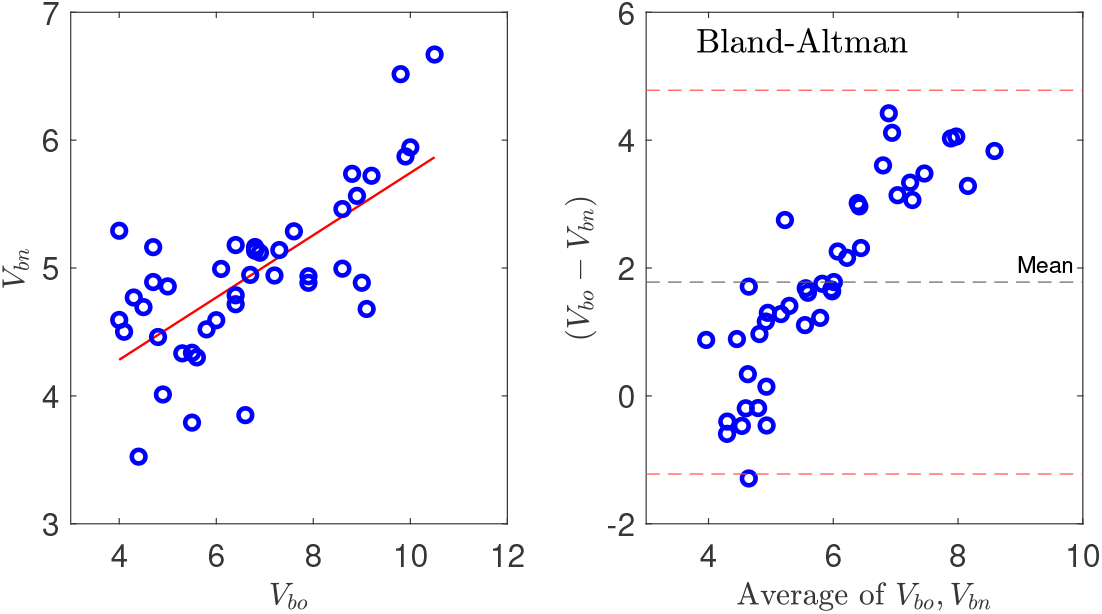
Comparison of methods. Left panel: Identity plot and linear regression (*V*_*bn*_ = 0.244 ∗ *V*_*bo*_ + 3.308, *r* = 0.5, red line) of blood volumes determined by previous (*V*_*bo*_) and new methods (*V*_*bn*_). Right panel: Bland-Altman analysis of difference of estimated volumes (*V*_*bn*_-*V*_*bo*_) versus average of both volumes, and mean of difference (black dashed line) and mean*±*2SD, red dashed lines.

An example of a poorly-modeled hematocrit course and the associated kernel probability distributions are presented in Fig. 6. Contrary to the poor modeling indicated by an increasing deviation between modeled and measured hematocrit values for advanced treatment phases, the hematocrit is perfectly predicted by the UKF approach throughout the whole observation phase. The associated KPD distribution in blood and fluid volumes is wider and bimodal. Similar distributions and variations in blood and fluid volume estimations were identified in six out of 21 treatments (28%).

**Fig. 6.**
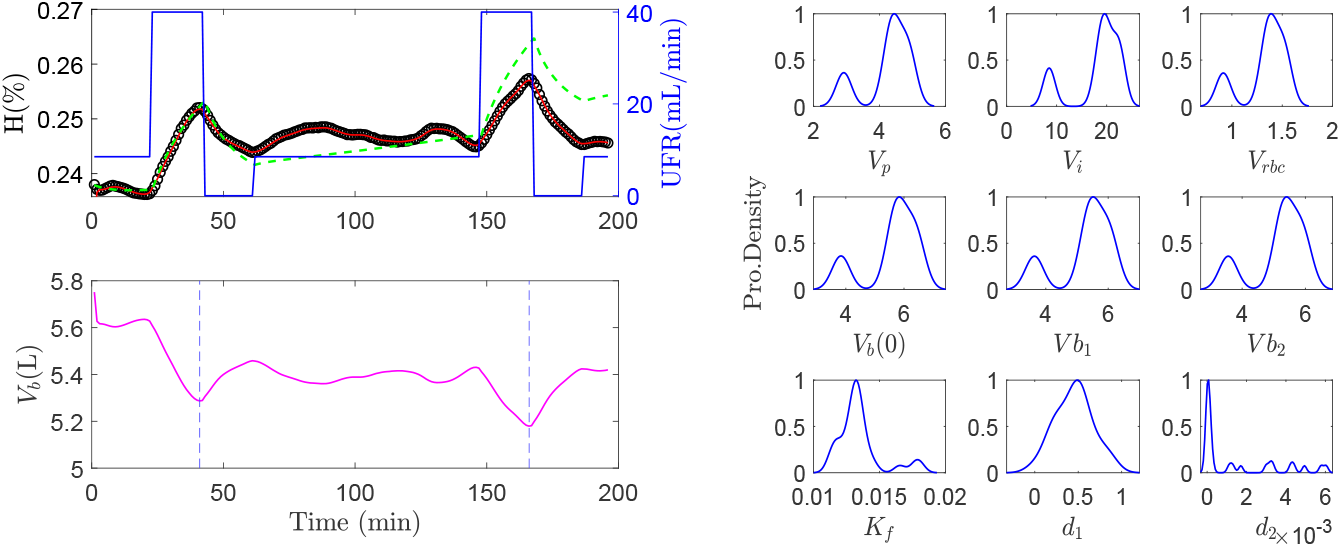
Modeling and prediction. Left, top panel: Poorly modeled hematocrit (green dashed trajectory) compared to predicted hematocrit (red trajectory) using UKF versus actual hematocrit (black circles) (data from treatment P82) in response to variable ultrafiltration rate (UFR, blue line). Left, bottom panel: Predicted blood volume (magenta) and time points (dashed blue lines) of blood volumes estimated at the end of ultrafiltration steps (*v*_1_, *v*_2_). Right panels: KPD functions for various parameters over the random initial solution Θ_0_ estimated for treatment P82.

## 4 Discussion

A new parameter and state estimation approach to identify blood volume in CKD patients is presented using hematocrit and ultrafiltration data continuously recorded during hemodialysis, a modified two-compartment volume kinetic model, anthropometric patient information, as well as specific treatment information. A short training phase was utilized to estimate the model parameters and initial conditions and then the rest of the data were used for model validation. Using this method, the whole time course of hematocrit was precisely predicted with a single set of estimated parameters, ultrafiltration rates, and the prediction (UKF) algorithm.

The experimental data show increased dynamics with two distinct hematocrit peaks within the first and the last hour of the 4 h observation phase (Fig. 3, Fig. 4) because ultrafiltration was varied between very high and very low rates for a different purpose. Such an ultrafiltration profile is atypical in normal dialysis prescription as ultrafiltration is usually delivered at a constant rate causing a much smoother hematocrit time course [20]. However, data recorded with variable ultrafiltration rates stimulating a characteristic system response have been used since the beginning of blood volume modeling [9]. In general they are better suited to test algorithms as parameter estimation and hematocrit prediction is more challenging with increased dynamics. The data are also characterized by low hematocrit values in the range of 25% as the measurements were done before erythropoiesis stimulating agents became available which typically increased hematocrit to about 35% [21].

In all treatments the predicted hematocrit perfectly fits the actual hematocrit (Fig. 3, Fig. 4, Fig. 6) whereas in six treatments the model alone (without UKF) failed to adequately describe the whole *H* profile. A closer look at one of these treatments (Fig. 6) reveals a discrepancy between modeled and measured hematocrit values starting in the period between ultrafiltration pulses. In this phase patients were allowed to sit up or to have a some small snack, all of which affects the driving forces for microvascular fluid shifts as well as the distribution of red blood cells within the cardiovascular space [22, 23]. As a result measured hematocrit varies independently of actual ultrafiltration rate. It is therefore no surprise that the effect of unknown inputs cannot adequately be modeled. Similar difficulties have previously been addressed by introducing an active control of lymphatic refilling [9].

The hematocrit prediction using the UKF algorithm is, however, perfect also in this case. In other words, even if the model is inadequate to describe the dynamics of the variable of interest, the UKF approach accounts for the error in model structure and still provides a perfect prediction.

This, however, introduces a different problem. Usually, the quality of a model is judged by the goodness of fit between modeled and measured data. The perfect fit between predicted (not modeled) and measured data using the UKF approach might be mistaken for a high quality of the model and model parameters and could be misleading. It is therefore important to also examine the precision of identified model parameters. In this study the KPD function for *V*_*b*_(0) is wide and bimodal (Fig. 6) and the resulting CV for initial blood volume is 16.3% and higher compared to the average CV of 9.8% for the whole group. The volume estimate in this specific treatment is therefore less precise, in spite of perfect hematocrit prediction using the UKF approach.

The value of the new approach is not so much in providing a precise prediction of the hematocrit, but in providing a measure for the precision of identified parameters. A precise prediction of hematocrit is not required because this variable is available and continuously measured by various on-line techniques. The practical interest of modeling is with precise identification of system parameters and variables which have clinical relevance but which cannot directly be measured. Parameters such as absolute blood volume must be inferred from modeling, usually by some type of indicator dilution. Different methods have been proposed and applied in clinical practice, but the precision of parameter identification is rarely addressed. Without information on precision, parameter identification is incomplete.

In this study the absolute blood volume was identified with an average CV of 9.8% (range 1 to 22%) while a precision of better than 5% would be desirable. An improved precision could be obtained by modifying and expanding the model, for example with regard to hemodynamic effects of ultrafiltration [24]. This, however requires more patient and treatment information and model assumptions, and it remains to be decided, whether such an expansion is feasible.

The method presented in this study can be used to identify critical specific blood volumes below which patients are more likely to experience intradialytic morbid events [25]. Specific blood volumes fell below the critical threshold of 65 mL/kg in three studies and were below that value from the beginning of treatment in one study. None of the patients became symptomatic during any of the treatments.

The proposed method could therefore assist in the prevention of IDH. Using the new approach, one could predict the risk of critical specific blood volumes ahead of time. This method could also be coupled to available feedback-control systems by providing the threshold for a critical blood volume but without relying on specific ultrafiltration profiles or volume infusions [26].

It is a limitation of the study that system parameters were not validated by accepted and independent reference methods. Absolute blood volumes were, however, compared to blood volumes identified by a different method applied to the same data set. This comparison provides a significant albeit moderate correlation, where volumes identified by the previous method appear to be increasingly overestimated at increasing blood volume range (Fig. 5). Extra-cellular volume modeled from whole-body bioimpedance spectroscopy could have been used to compare the sum of plasma and interstitial volumes. It also remains to be tested whether the new approach provides precise and non-trivial volume estimates in treatments with constant ultrafiltration rates.

## 5 Conclusions

The new estimation approach offers a practical method for estimating absolute blood volume in everyday dialysis without relying on a specific ultrafiltration or volume infusion protocol. The absolute blood volume estimated at the beginning and during every dialysis session offers the opportunity to improve fluid management in CKD patients. This method also can be used to determine critical specific blood volumes during dialysis treatment and to detect low absolute blood volumes ahead of time. The immediate impact of this work is the development of a new approach for estimating absolute blood volume during dialysis sessions which is ready to be implemented into dialysis machines used for treating both chronic and acute renal failure.

## Supporting information

Results, model, and tables

## Acknowledgements

The authors would like to thank Prof. Yossi Chait, University of Massachusetts Amherst, for his help and support.

## Authors’ Biographies

**Rammah Abohtyra:** received his Ph.D. degrees in electrical engineering from Colorado School of Mines Golden, CO, USA, in 2015. As a Postdoc Scholar at the Pennsylvania State University, University Park, PA, USA, his primary research focuses on leveraging biological modeling and control system to improve health problems, with different medical applications.

**Tyrone Vincent:** received his Ph.D. degrees in electrical engineering from the University of Michigan, Ann Arbor, MI, USA, in 1997. He is currently a Professor in the Department of Electrical Engineering and Computer Science at the Colorado School of Mines, Golden, CO, USA. His research interests include system identification, estimation, and control with different applications.

**Daniel Schneditz** received his Ph.D. degree in chemistry and experimental physics from the Karl-Franzens University, Graz, Austria in 1983. He is currently affiliated with the Department of Physiology from the Medical University in Graz, Austria from which he retired as an Associated Professor in 2021. His research is focused on cardiovascular, fluid and electrolyte physiology, extracorporeal sensors, and mathematical modeling with emphasis on mechanistic, physiology-based modeling in applications with extracorporeal systems such as hemodialysis for renal replacement therapy.

