## Supplementary material for "Nonlinear Parameter and State Estimation Approach for Intradialytic Measurement of Absolute Blood Volume": Results, model, and tables

### Supplementary File

### Blood volume model

The microvascular (shift/refill) filtration and lymphatic flows are given by:

$$Q_f(V_{pl}, V_{int}) = K_f(\delta_{pl} - \delta_\pi) \tag{1}$$
$$Q_l(V_{int}) = g\tanh(h\pi_{int}) + \beta \tag{2}$$

where $\delta_{pl}$ and $\delta_\pi$ represent the hydrostatic and colloid osmotic pressure gradients, respectively:

$$\delta_{pl}(t) = (p_c(t) - p_{int}(t)) \tag{3}$$
$$\delta_\pi(t) = (\pi_{pl} - \pi_{int}(t)) \tag{4}$$

and

$$p_c(t) = d_1(100(\frac{V_{rbc} + V_{pl}(t)}{V_{rbc} + V_{p,eu}}) + r)^{d_2} + p_0 \tag{5}$$
$$p_{int}(t) = (100(\frac{\mu V_{int}(t)}{V_{int,eu}}) + \frac{\lambda}{(100\frac{V_{int}}{V_{int,eu}} + \gamma)}) \tag{6}$$
$$\pi_{pl}(t) = (\frac{k_{c1}M_{pl}}{V_{pl}(t)} + \frac{k_{c2}M_{pl}^2}{V_{pl}^2(t)} + \frac{k_{c3}M_{pl}^3}{V_{pl}^3(t)}) \tag{7}$$
$$\pi_{int}(t) = (\frac{k_{c1}M_{int}}{V_{int}(t)} + \frac{k_{c2}M_{int}^2}{V_{int}^2(t)} + \frac{k_{c3}M_{int}^3}{V_{int}^3(t)}) \tag{8}$$

where $p_c$, $p_{int}$, $\pi_{pl}$, $\pi_{int}$ refer to the hydrostatic capillary pressure, interstitial pressure, plasma colloid osmotic pressure, and interstitial colloid osmotic pressure, respectively, and $g$, $h$, and $\beta$ are constants; and where $V_{p,eu}$, $V_{int,eu}$ denote euhydration (normal) plasma and interstitial volumes for a 70 kg patient, respectively;$p_0$ is an offset pressure; $M_{pl}$ is plasma protein mass; $M_{int}$ is interstitial protein mass, all are listed in Table 1, in which $\mu$, $\lambda$, $\gamma$, $d_1$, $d_2$, $r$, $M_{pl}$, $M_{int}$, $k_{c1}$, $k_{c2}$ and $k_{c3}$ are nominal parameters of model (1).

| Constants | value |
| --- | --- |
| $\mu$ | 0.006 |
| $\lambda$ | -198 |
| $\gamma$ | -45 |
| $g$ | 0.045 |
| $h$ | 0.767 |
| $\beta$ | 0.045 |
| $p_0$(mmHg) | 13.128 |
| $V_{p,eu}$ (L) | 3 |
| $V_{int,eu}$ (L) | 11 |
| $k_{c1}$ | 0.21 |
| $k_{c2}$ | 0.0016 |
| $k_{c3}$ | 9e-6 |

Table 1: The nominal parameters of the nonlinear two-compartment model.

### Patient and treatment information

A patient-specific and treatment information is provided in this table.

| Patient ID | Sex | BW pre [kg] | BW post [kg] | UFV [L] | time [min] | $H(0)$ [%] | $c_p(0)$ [g/L] |
|---|---|---|---|---|---|---|---|
| P11 | m | 76.2 | 74.0 | 2.5 | 240 | 28.3 | 66 |
| P12 | m | 76.5 | 74.0 | 2.8 | 270 | 29.1 | 70.4 |
| P21 | f | 62.8 | 60.0 | 3.3 | 240 | 19.0 | 58.7 |
| P22 | f | 63.2 | 60.0 | 3.2 | 240 | 18.1 | 65.2 |
| P23 | f | 62.5 | 60.0 | 3.0 | 240 | 17.8 | 63.9 |
| P31 | f | 52.9 | 50.0 | 3.7 | 360 | 26.2 | 64.3 |
| P32 | f | 53.3 | 50.0 | 4.1 | 360 | 24.8 | 63.6 |
| P33 | f | 52.5 | 50.0 | 2.7 | 360 | 24.7 | 65.3 |
| P41 | m | 67.8 | 64.5 | 3.6 | 270 | 25.5 | 72.5 |
| P42 | m | 68.5 | 64.8 | 4.3 | 240 | 25.9 | 74.9 |
| P43 | m | 67.1 | 64.5 | 2.9 | 240 | 25.0 | 70.8 |
| P51 | m | 62.3 | 58.0 | 4.9 | 270 | 20.7 | 68.8 |
| P52 | m | 61.2 | 58.0 | 3.8 | 240 | 19.1 | 63.4 |
| P61 | f | 56.4 | 52.8 | 4.7 | 270 | 25.0 | 64.2 |
| P62 | f | 57.0 | 52.5 | 4.8 | 270 | 26.4 | 63.1 |
| P71 | m | 117.3 | 110.0 | 7.6 | 420 | 22.1 | 67.6 |
| P81 | m | 67.1 | 64.0 | 3.7 | 480 | 23.2 | 65.9 |
| P82 | m | 68.4 | 64.0 | 5.0 | 480 | 23.8 | 69.3 |
| P83 | m | 68.3 | 64.0 | 4.9 | 480 | 24.1 | 67.8 |
| P91 | f | 53.5 | 50.5 | 3.3 | 240 | 20.3 | 70.1 |
| P101 | m | 62.2 | 58.5 | 5.0 | 300 | 21.6 | 68.4 |
| mean | | 65.57 | 62.10 | 3.99 | 324.3 | 23.37 | 66.87 |
| SD | | 13.44 | 12.76 | 1.14 | 85.52 | 1.65 | 3.66 |

Table 2: Patient information. Sex: male (m), female (f), pre-dialysis body weight (BWpre), post-dialysis body wight (BW post), ultrafiltration volume (UFV), treatment time,initial hematocrit $H(0)$ (%), and initial plasma protein concentration $C_p(0)$.

#### Estimated parameters

Parameter estimation results (mean±SD) using the algorithm for 21 treatments.

| Patient ID Number | $K_f$ (L/(min*mmHg)) | $d_1$ (-) | $d_2$ (-) | $V_{rbc}$ (L) |
|---|---|---|---|---|
| P11 | 0.0053±6e-4 | 0.8±0.44 | 4.1e-3±3e-3 | 1.7±0.07 |
| P12 | 0.0059±4e-4 | 0.9±0.34 | 6.1e-3±4e-3 | 1.6±0.02 |
| P21 | 0.0143±3e-3 | 0.9±0.19 | 9.7e-3±6e-3 | 0.9±0.09 |
| P22 | 0.0246±3e-3 | 0.3±0.26 | 5e-4±1e-3 | 0.9±0.09 |
| P23 | 0.0209±3e-3 | 0.5±0.14 | 1.2e-3±2e-3 | 0.9±0.05 |
| P31 | 0.0086±3e-3 | 1.2±0.17 | 1e-2±5e-3 | 1.3±0.09 |
| P32 | 0.0099±3e-3 | 1.1±0.24 | 9.6e-3±5e-3 | 1.3±0.14 |
| P33 | 0.0127±3e-3 | 0.6±0.33 | 2.3e-3±3e-3 | 1.4±0.21 |
| P41 | 0.0841±3e-3 | 0.9±0.39 | 5.6e-3±4e-3 | 1.3±0.16 |
| P42 | 0.0171±3e-3 | 0.33±0.29 | 6e-4±1e-3 | 1.7±0.10 |
| P43 | 0.0343±3e-3 | 0.9±0.19 | 7.8e-3±3e-3 | 1.5±0.05 |
| P51 | 0.0253±3e-3 | 0.2±0.19 | 5e-4±2e-3 | 1.2±0.03 |
| P52 | 0.0256±3e-3 | 0.4±0.20 | 4e-4±1e-3 | 1.0±0.05 |
| P61 | 0.0093±3e-3 | 0.6±0.32 | 0.0028±3e-3 | 1.4±0.21 |
| P62 | 0.0065±3e-3 | 0.9±0.38 | 0.0054±3e-3 | 1.3±0.21 |
| P71 | 0.0193±3e-3 | 0.6±0.24 | 0.0023±2e-3 | 1.1±0.14 |
| P81 | 0.0110±3e-3 | 0.9±0.22 | 0.0073±6e-3 | 1.2±0.17 |
| P82 | 0.0135±3e-3 | 0.5±0.45 | 0.0015±2e-3 | 1.3±0.22 |
| P83 | 0.0176±3e-3 | 0.3±0.29 | 0.0019±2e-3 | 1.3±0.27 |
| P91 | 0.0107±3e-3 | 0.9±0.23 | 0.0116±6e-3 | 0.9±0.07 |
| P101 | 0.0048±3e-3 | 1.1±0.26 | 0.0231±1.5e-3 | 0.9±0.13 |
| Average | 0..0182±4.5e-3 | 0.7±0.25 | 0.0054±3.7e-3 | 1.3±0.12 |

Table 3: Model parameter estimation results (mean±SD) using the algorithm for 21 treatments, including the filtration coefficient ($K_f$), and the model hydrostatic pressure parameters ($d_1$ and $d_2$), and red blood volume.
